# Molecular dissection of early defense signaling underlying volatile-mediated defense priming and herbivore resistance in rice

**DOI:** 10.1101/378752

**Authors:** Meng Ye, Gaétan Glauser, Yonggen Lou, Matthias Erb, Lingfei Hu

## Abstract

Herbivore-induced plant volatiles prime plant defenses and resistance. How volatiles are integrated into early defense signaling is not well understood. Furthermore, whether there is a causal relationship between volatile defense priming and herbivore resistance is unclear. Here, we investigated the impact of indole, a common herbivore-induced plant volatile and known defense priming cue, on early defense signaling and herbivore resistance in rice. We show that rice plants infested by *Spodoptera frugiperda* caterpillars release up to 25 ng*h^−1^. Exposure to equal doses of synthetic indole enhances rice resistance to *S. frugiperda*. Screening of early signaling components reveals that indole directly enhances the expression of the receptor like kinase *OsLRR-RLK1*. Furthermore, indole specifically primes the transcription, accumulation and activation of the mitogen-activated protein kinase *OsMPK3* as well as the expression of the downstream WRKY transcription factor *OsWRKY70* and several jasmonate biosynthesis genes, resulting in a higher accumulation of jasmonic acid (JA). Using transgenic plants defective in early signaling, we show that *OsMPK3* is required, and that *OsMPK6* and *OsWRKY70* contribute to indole-mediated defense priming of JA-dependent herbivore resistance. We conclude that volatiles can increase herbivore resistance of plants by priming early defense signaling components.

## Introduction

Plants that are under attack by insect herbivores emit specific blends of herbivore-induced plant volatiles (HIPVs). HIPVs can prime intact plant tissues to respond faster and/or stronger to subsequent herbivore attack (Ton et al., 2007; Kim and Felton, 2013; Balmer et al., 2015; Erb et al., 2015; Mauch-Mani et al., 2017) and may thereby act as within-plant defense signals that overcome vascular constraints (Frost et al., 2007; Heil and Silva Bueno, 2007).

HIPVs can prime plants to the accumulation of jasmonate defense regulators. Maize (Zea *mays)* HIPVs such as indole prime jasmonic acid (JA) accumulation and the transcription of jasmonate-responsive genes (Ton et al., 2007; Erb et al., 2015). Similarly, green leaf volatiles (GLVs) such as (Z)-3-hexenyl acetate prime JA production in maize (Engelberth et al., 2004) and hybrid poplar *(Populus deltoides* × *nigra*) (Frost et al., 2008). As jasmonates are important regulators of plant defense and herbivore resistance (Howe and Jander, 2008), and several HIPVs prime jasmonates, it is generally assumed that HIPVs increase plant resistance by priming the jasmonate pathway (Engelberth et al., 2004; Ameye et al., 2015). However, this connection has never been directly tested. Recent work shows that some HIPVs can also increase plant resistance directly by being absorbed and transformed into toxins (Sugimoto et al., 2014). Thus, the relative importance of HIPV-mediated defense priming for herbivore resistance remains unclear.

To date, several HIPV priming cues have been identified, and their impact on early defense signaling has been investigated (Shulaev et al., 1997; Engelberth et al., 2013; Erb et al., 2015). In maize, (Z)-3-hexenol increases the expression of *WRKY12* and *MAPK6*, which are likely involved in transcriptional defense regulation. The same volatile also activates putative JA biosynthesis genes such as *AOS* and *LOX5* (Engelberth et al., 2013). In *Arabidopsis thaliana*, (E)-2-hexenal induces the expression of *WRKY40* and *WRKY6* (Mirabella et al., 2015). *WRKY40* and *WRKY6* regulate γ-amino butyric acid (GABA) metabolism, which mediates GLV-induced root growth suppression in a JA-independent manner (Mirabella et al., 2008). Despite these promising results, how HIPVs are integrated into early defense signaling to regulate JA-dependent defenses remains unclear.

We recently identified indole as an herbivore-induced volatile within-plant signal that primes JA and is required for the systemic priming of monoterpenes in maize (Erb et al., 2015). Indole also primes volatiles in cotton, suggesting that it is active across different plant species (Erb et al., 2015). Indole can furthermore interact with (Z)-3-hexenyl acetate to increase JA signaling and herbivore resistance (Hu et al., under review). Indole exposure also directly increases the mortality of early instar *Spodoptera littoralis* caterpillars by approx. 10% (Veyrat et al., 2016) and renders caterpillars more resistant and less attractive to parasitoids (Ye et al., 2018). To understand if and how indole is integrated into early defense signaling, we studied its role on rice. Rice is a useful model, as several key players in early defense signaling have been identified, including receptor-like kinases (Ye, 2016; Hu et al., 2018), mitogen-activated protein kinase (MPKs) (Wang et al., 2013; Li et al., 2015; Liu et al., 2018), WRKY transcription factors (Wang et al., 2007; Hu et al., 2015; Li et al., 2015; Hu et al., 2016; Huangfu et al., 2016) and jasmonate biosynthesis genes (Zhou et al., 2009; Guo et al., 2014; Hu et al., 2015). By taking advantage of the available knowledge and molecular resources in rice, we investigated how indole is integrated into early defense signaling, and to what extent this integration translates into enhanced herbivore resistance.

## Results

### Caterpillar-induced indole increases herbivore resistance

To determine whether caterpillar attack induces the release of indole in rice, we infested rice plants with *Spodoptera frugiperda* caterpillars and measured indole release rates 12 - 20 h after the beginning of the attack. Indole emissions increased with the severity of *S. frugiperda* attack and ranged from 9 to 25 ng h^−1^ per plant (Figure 1A-C). Based on these results, we calibrated capillary dispensers to release indole at a physiologically relevant rate of 21 ng h^−1^ (Figure 1C) and exposed rice plants to individual dispensers for 12 h. We then added *S. frugiperda* larvae to control and indole-exposed plants and measured larval weight gain and plant damage. Indole pre-exposure significantly reduced larval damage and weight gain (Figure 1D, E). Thus, physiologically relevant concentrations of indole are sufficient to increase rice resistance against a chewing herbivore.

**Figure 1.**
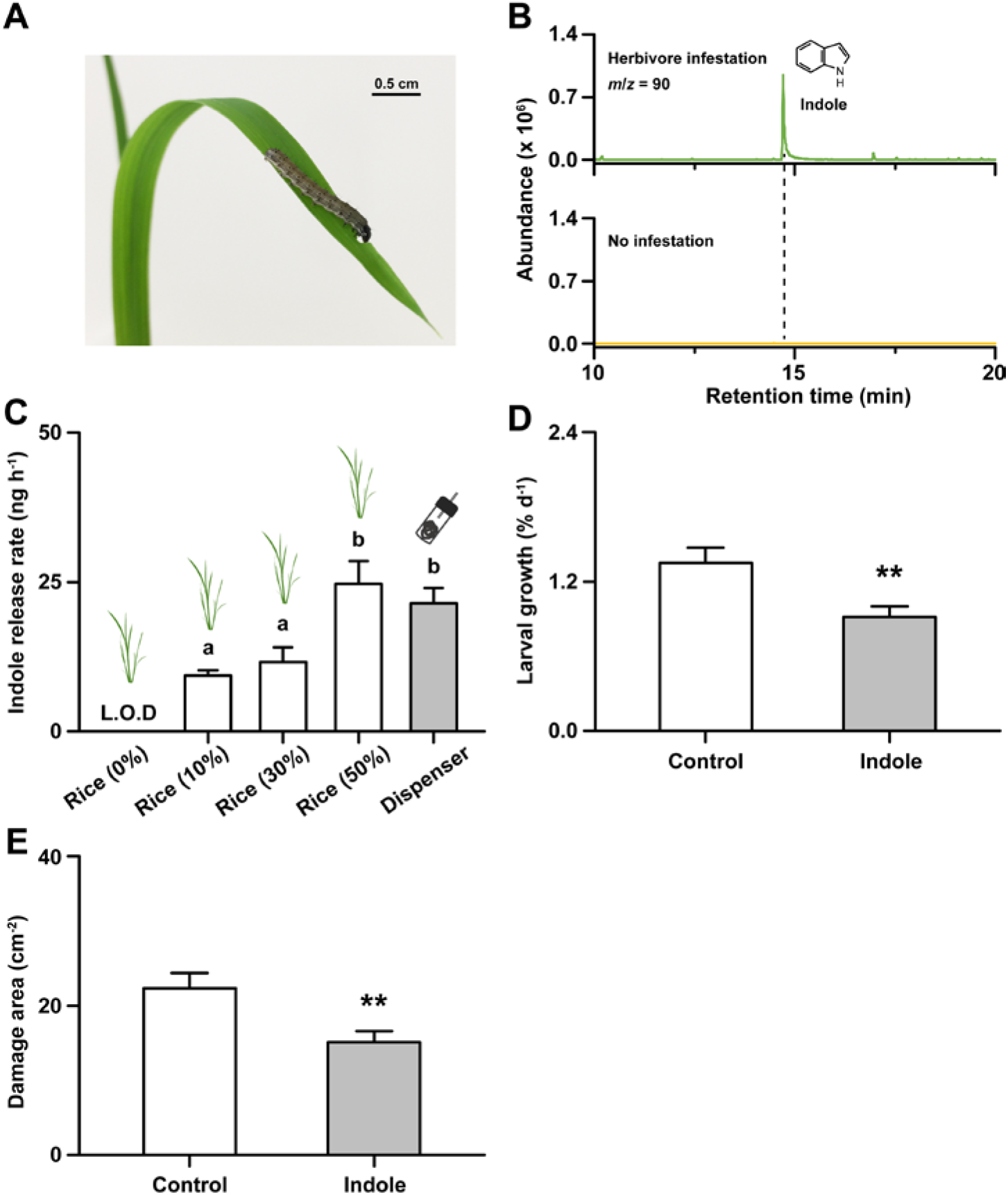
Indole is an herbivore-induced plant volatile that increases rice resistance to *Spodoptera frugiperda* larvae at physiological doses. (**A**) An *S. frugiperda* caterpillar feeding on a rice leaf. (**B**) Extracted ion chromatograms of GC/MS headspace analyses of control and *S. frugiperda* infested rice leaves. *m/z* = 90 corresponds to a characteristic fragment of indole. (**C**) Emission rates of indole from rice plants that are attacked by different densities of *S. frugiperda* caterpillars. The percentage of consumed leaf area relative to total leaf area is indicated on the x-axis (+SE, *n*=6-8). The release of synthetic indole by custom-made capillary dispensers is shown for comparison. Letters indicate significant differences between treatments (*P* < 0.05, one-way ANOVA followed by multiple comparisons through FDR-corrected LSMeans). L.O.D., below limit of detection. (**D**) Average growth rate of *S. frugiperda* caterpillars feeding on rice plants that were pre-exposed to indole dispensers releasing indole at approx. 21 ng h^−1^ or control dispensers for 12 h prior to infestation (+SE, *n*=15). (**E**) Average consumed leaf area (+SE, *n*=15). Asterisks indicate significant differences between the volatile exposure treatments (Student’s *t*-tests, **, *P* < 0.01).

### Indole primes the transcription of early defense signaling genes

To explore the capacity of indole to regulate early defense signaling in rice, we profiled the expression of known early defense signaling genes (Figure 2A), including two receptor-like kinases (Ye, 2016; Hu et al., 2018), two MPKs (Wang et al., 2013; Li et al., 2015), seven WRKY transcription factors (Qiu et al., 2008; Koo et al., 2009; Li, 2012; Han et al., 2013; Hu et al., 2015; Li et al., 2015; Huangfu et al., 2016) and five jasmonate biosynthesis genes (Figure 2A) (Zhou et al., 2009; Fukumoto et al., 2013; Guo et al., 2014; Hu et al., 2015). Control plants and plants that were pre-exposed to indole for 12 h were measured 0 min, 90 min and 360 min after simulated herbivore attack to capture both direct induction and priming. Herbivory was simulated by wounding the leaves and adding *S. frugiperda* oral secretions (OS) as described (Erb et al., 2009; Fukumoto et al., 2013; Chuang et al., 2014). The expression of *OsLRR-RLK1*, a receptor like kinase that regulates herbivore resistance (Hu et al., 2018), was directly induced by indole exposure and expressed at higher levels 90 minutes after simulated herbivore attack (Figure 2B). The transcription of *OsMPK3*, an MPK which acts downstream of *OsLRR-RLK1* to regulate herbivore-induced defense and resistance (Wang et al., 2013; Hu et al., 2018) was not directly induced by indole, but primed for higher expression 90 min after simulated *S. frugiperda* attack (Figure 2C). *OsWRKY70*, which is a positive regulator of herbivore-induced defense and acts downstream of *OsMPK3* (Li et al., 2015), was primed in a similar manner (Figure 2D). Three jasmonate biosynthesis genes, *OsAOS1, OsAOC* and *OsOPR3* were equally primed by indole 90 minutes after elicitation (Figure 2E). By contrast, *OsHI-RLK2, OsMPK6, OsWRKY13, OsWRKY24, OsWRKY30, OsWRKY33, OsWRKY45, OsWRKY53* and the jasmonate biosynthesis genes *OsHI-LOX* and *OsJAR1* did not respond to indole pretreatment (Figure 2B-E). Thus, indole increases the expression of a specific subset of early defense signaling genes upstream of JA biosynthesis.

**Figure 2.**
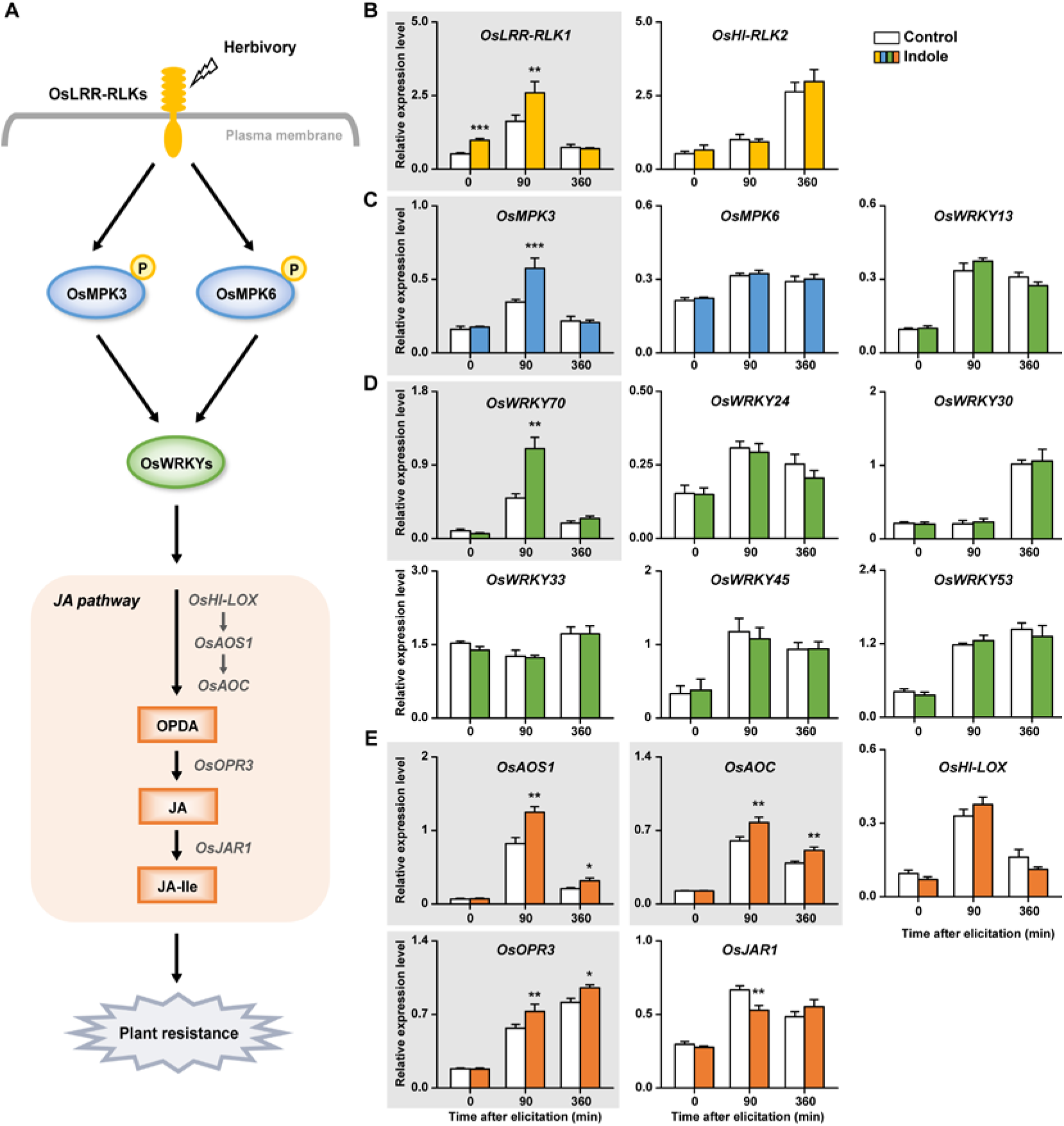
Indole primes early defense signaling genes. (**A**) Current model of herbivory-induced defense signaling in rice, including leucine-rich repeat receptor-like kinases (LRR-RLKs), mitogen-activated protein kinases (MPKs), WRKY transcription factors, jasmonate biosynthesis genes and oxylipins. (**B – E**) Effect of indole pre-treatment on the expression of genes coding for the different early signaling steps at different time points after elicitation by wounding and application of *Spodoptera frugiperda* oral secretions (+SE, *n*=4-6). OPDA, 12-oxophytodienoic acid; JA, jasmonic acid; JA-Ile, JA-isoleucine. Asterisks indicate significant differences between volatile exposure treatments at different time points (two-way ANOVA followed by pairwise comparisons through FDR-corrected LSMeans; *, *P* < 0.05; **, *P* < 0.01; ***, *P* < 0.001). Genes responding to indole are highlighted in gray.

### Indole primes OsMPK3 accumulation and activation

To determine whether transcriptional priming of MPKs is also reflected in protein abundance, we performed western blots using OsMPK3 and OsMPK6-specific antibodies. Protein accumulation of OsMPK3 was primed by indole, leading to higher OsMPK3 abundance 90 min after elicitation (Figure 3A). OsMPK6 accumulation was not altered by indole pre-treatment (Figure 3B). To further investigate whether indole pretreatment increases OsMPK3 activation, we measured OsMPK3 phosphorylation by immunoblot analysis using an anti-phosphoERK1/2 (anti-pTEpY) antibody that interacts with doubly phosphorylated (activated) MPK3 and MPK6 (Segui-Simarro et al., 2005; Anderson et al., 2011; Schwessinger et al., 2015). Indole primed OsMPK3 activation 90 min after elicitation (Figure 3C). We also detected a slightly higher activation of OsMPK6 (Figure 3C). Thus, indole exposures primes the accumulation and activation of MPKs involved in defense regulation.

**Figure 3.**
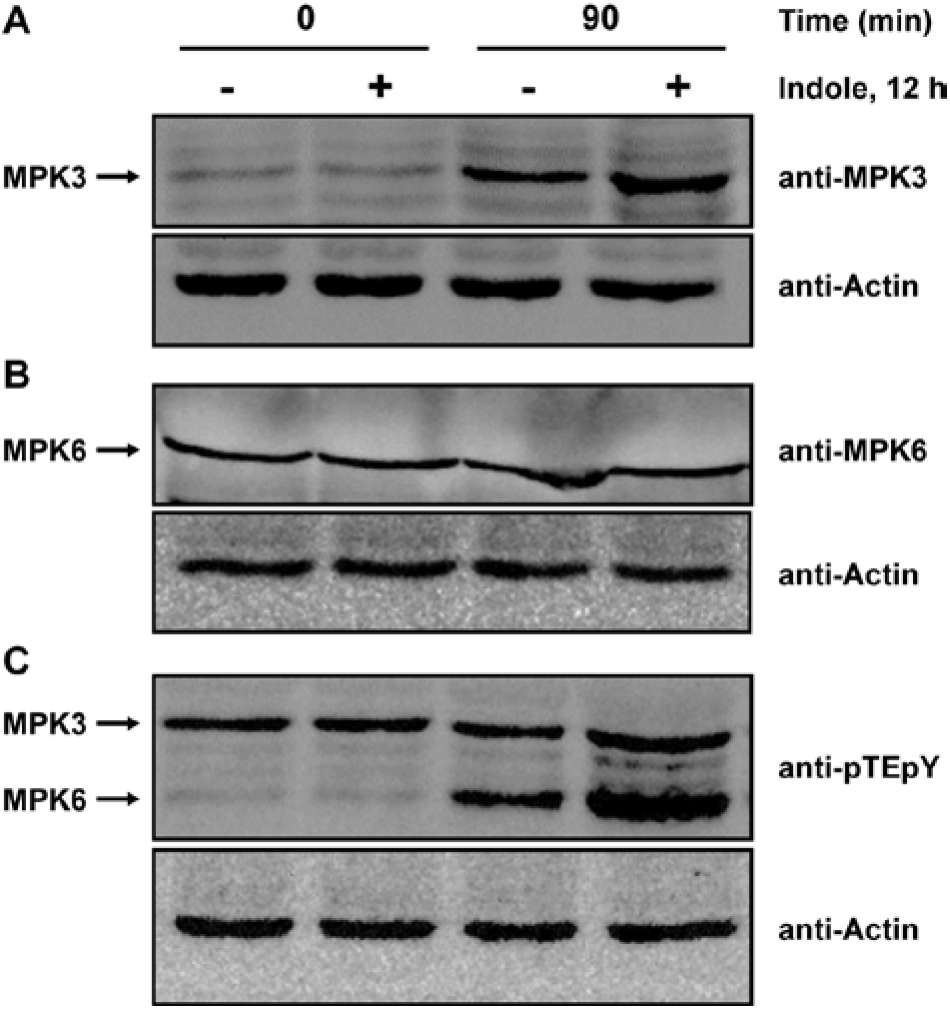
Indole primes OsMPK3 accumulation and activation. (**A – C**) Herbivore-elicited protein accumulation and activation of OsMPK3 and OsMPK6 with (+) or without (-) indole exposure for 12 h. Leaves from 6 replicate plants were harvested at indicated times after elicitation. Immunoblotting was performed using an anti-MPK3 antibody for OsMPK3 (A), an anti-MPK6 antibody to for OsMPK6 (B), an anti-pTEpY antibody to detect phosphorylated MPKs (C), or an actin antibody as a loading control. Actin was measured on a replicate blot. This experiment was repeated two times with similar results.

### Indole induces OPDA and primes JA

To investigate whether the activation of early defense signaling components is associated with higher accumulation of stress-related phytohormones, we quantified 12-oxophytodienoic acid (OPDA), JA and JA-isoleucine (JA-Ile), abscisic acid (ABA) and salicylic acid (SA) in indole-exposed and control plants (Figure 4).

**Figure 4.**
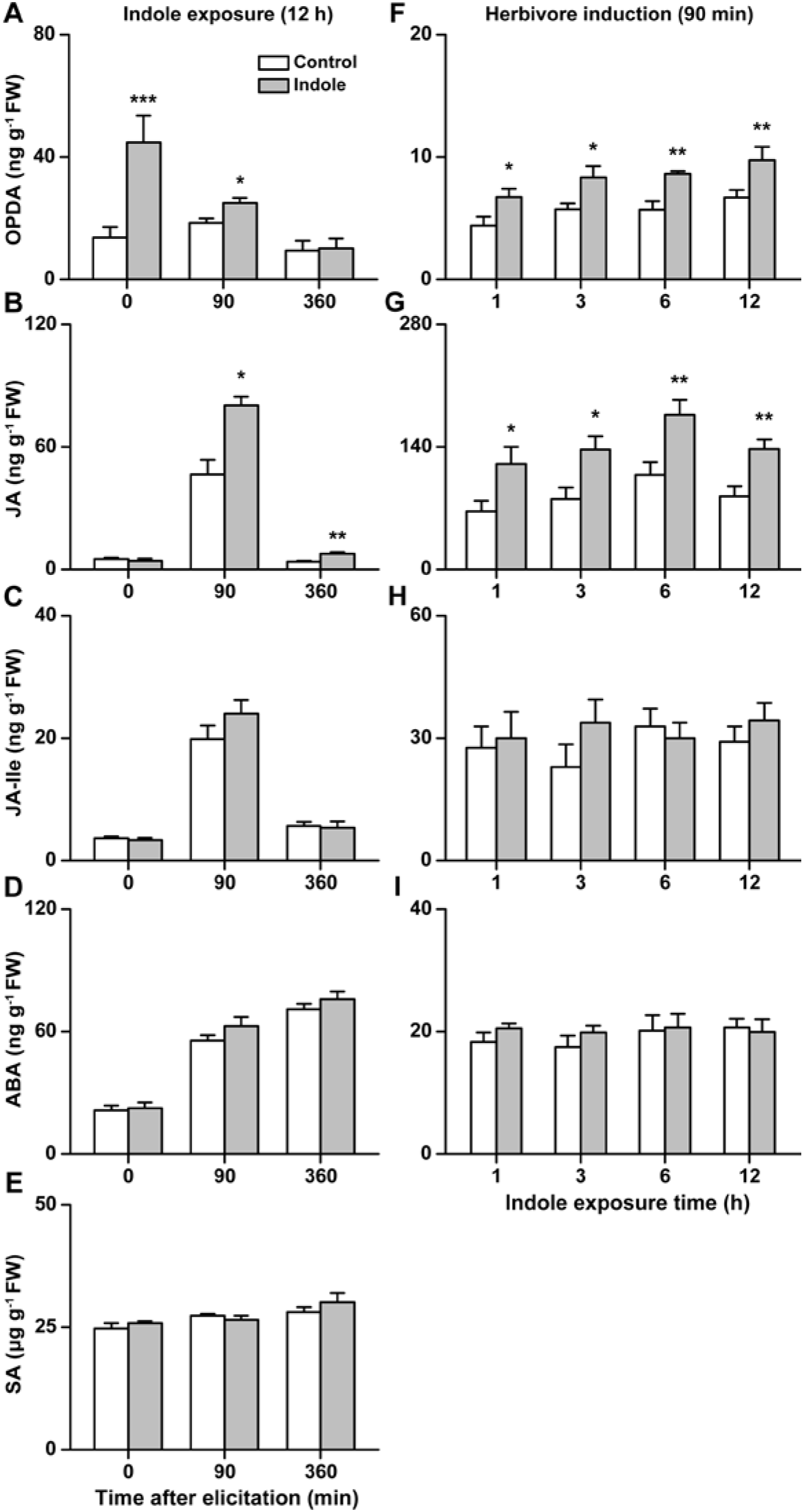
Indole induces 12-oxophytodienoic acid (OPDA) and primes jasmonic acid (JA) accumulation. (**A – E**) Average concentrations of (**A**) OPDA, (**B**) JA, (**C**) JA-isoleucine (JA-Ile), (**D**) abscisic acid (ABA) and (**E**) salicylic acid (SA) in indole- and control-exposed rice plants at different time points after elicitation (+SE, *n*=5-6). Plants were exposed to indole for 12 h before elicitation. (**F – I**) Average concentrations of OPDA, JA, JA-Ile and ABA in rice plants that were exposed to indole for 1 h, 3 h, 6 h or 12 h or control dispensers 90 min after elicitation (+SE, *n*=5-6). SA levels were not measured in this experiment. Asterisks indicate significant differences between treatments (two-way ANOVA followed by pairwise comparisons through FDR-corrected LSMeans; *, *P* < 0.05; **, *P* < 0.01; ***, *P* < 0.001).

Indole exposure increased the accumulation of OPDA before and after elicitation (Figure 4A). JA concentrations were increased in indole-exposed plants 90 and 360 min after elicitation (Figure 4B). The levels of JA-Ile, SA and ABA were not affected by indole pre-exposure (Figure 4C-E). To test the total dose of indole that is required for the priming of phytohormones, we exposed rice plants to indole dispensers for 1-12 h and measured hormone accumulation 90 minutes after elicitation. Exposure to indole dispensers for 1 h (resulting in a total release of 21 ng from the dispensers) was sufficient to increase OPDA and JA levels. Longer exposure did not significantly increase OPDA and JA responses. Thus, exposure of rice plants to 21 ng of indole over 1 h is sufficient to increase the production of oxylipin defense regulators.

### OsMPK3 is required for indole-elicited JA priming and herbivore resistance

To understand whether the early signaling components that are responsive to indole are required for downstream responses, we measured JA priming and herbivore resistance in control- and indole-exposed wild type and transgenic plants, including the *OsLRR*-RLK1-silenced line *ir-lrr1* (Hu et al., 2018), the *OsMPK3*- and OsMPK6-silenced lines *ir-mpk3* and *ir-mpk6* (Wang et al., 2013; Li et al., 2015) and the OsWRKY70-silenced line *ir-wrky70* (Li et al., 2015). *OsLRR-RLK1* silencing did not affect indole-dependent OPDA induction, JA priming and herbivore growth suppression (Figure 5A). By contrast, silencing *OsMPK3* completely suppressed indole-dependent OPDA induction, JA priming and herbivore growth reduction (Figure 5B). The induction of JA by herbivore elicitation was still clearly visible in *ir-mpk3* plants, demonstrating that the absence of indole resistance priming is not due to a complete suppression of JA signaling. Silencing *OsMPK6* reduced indole-dependent JA priming and herbivore growth suppression by approximately 50%, and led to an almost complete disappearance of OPDA induction. Silencing *OsWRKY70* also reduced indole-dependent OPDA induction, priming of JA and herbivore growth suppression by approximately 50 % (Figure 5C, D). Thus, *OsMPK3* is required, and *OsMPK6* and *OsWRKY70* contribute to indole defense priming.

**Figure 5.**
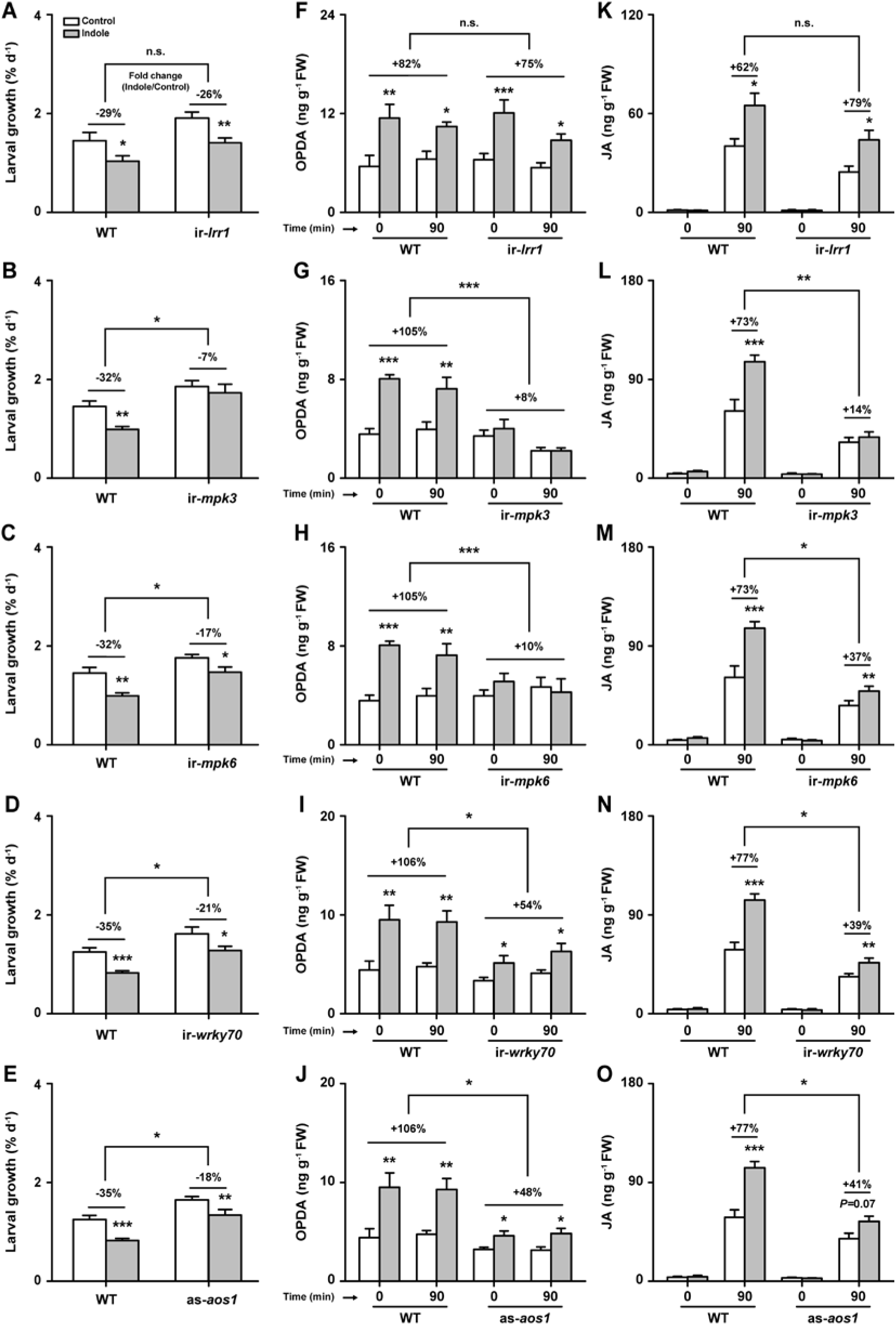
Indole-induced priming of jasmonic acid and herbivore resistance depends on OsMPK3. (**A – E**) Average growth rate of *Spodoptera frugiperda* caterpillars feeding on (**A**) *ir-lrr1*, (**B**) *ir-mpk3*, (**C**) *ir-mpk6*, (**D**) *ir-wrky70*, (**E**) *as-aos1* lines and wild-type (WT) plants that were preexposed to indole or control (+SE, *n*=15). (**F – J**) Average concentrations of herbivore-induced 12-oxophytodienoic acid (OPDA) in the different transgenic lines and WT plants that were pre-exposed to indole or control dispensers (+SE, *n*=6). (**K – O**) Average concentrations of herbivore-induced jasmonic acid (JA) in the different transgenic lines and WT plants that were pre-exposed to indole or control dispensers (+SE, *n*=6). Note that WT, *ir-mpk3* and *ir-mpk6* plants as well as WT, *ir-wrky70* and *as-aos1* plants were measured together within the same experiments. The WT data is, therefore, identical in the respective figures (e.g. same WT data for *ir-mpk3* and *ir-mpk6* figures; same WT data for *ir-wrky70* and *as-aos1* figures) and shown repeatedly for illustrative purposes. FW, fresh weight. n.s. not significant. Percentages refer to fold changes of indole-exposed plants relative to control-exposed plants. Asterisks above bars indicate significant differences between volatile exposure treatments within the same plant genotype (two-way ANOVA followed by pairwise comparisons through FDR-corrected LSMeans; *, *P* < 0.05; **, *P* < 0.01; ***, *P* < 0.001). Asterisks above bars represent significant differences between indole-dependent fold changes of WT and transgenic lines (Student’s *t*-tests, *, *P* < 0.05; **, *P* < 0.01; ***, *P* < 0.001).

### The jasmonate signaling pathway contributes to indole-induced herbivore resistance

To study the connection between the regulation of JA and the decrease in herbivore performance in indole-exposed plants, we tested *as-aos1* plants, which accumulate lower levels of jasmonates upon herbivore elicitation (Hu et al., 2015). OPDA, accumulation, JA priming and herbivore growth suppression were reduced by approximately 50% in *as-aos1* plants (Figure 5E). Across the different genotypes, herbivore growth suppression was strongly correlated with OPDA and JA overaccumulation: Genotypes that responded to indole with stronger OPDA accumulation and JA priming also reduced larval growth more strongly after pre-exposure (Figure 6A, B). By contrast, JA-Ile did not respond significantly to indole pre-treatment in any of the measured genotypes (Supplemental Figure 1), and there was no correlation between indole-effects on JA-Ile and herbivore growth suppression (Figure 6C). Together, these findings implicate the jasmonate signaling pathway in indole-induced herbivore resistance.

**Figure 6.**
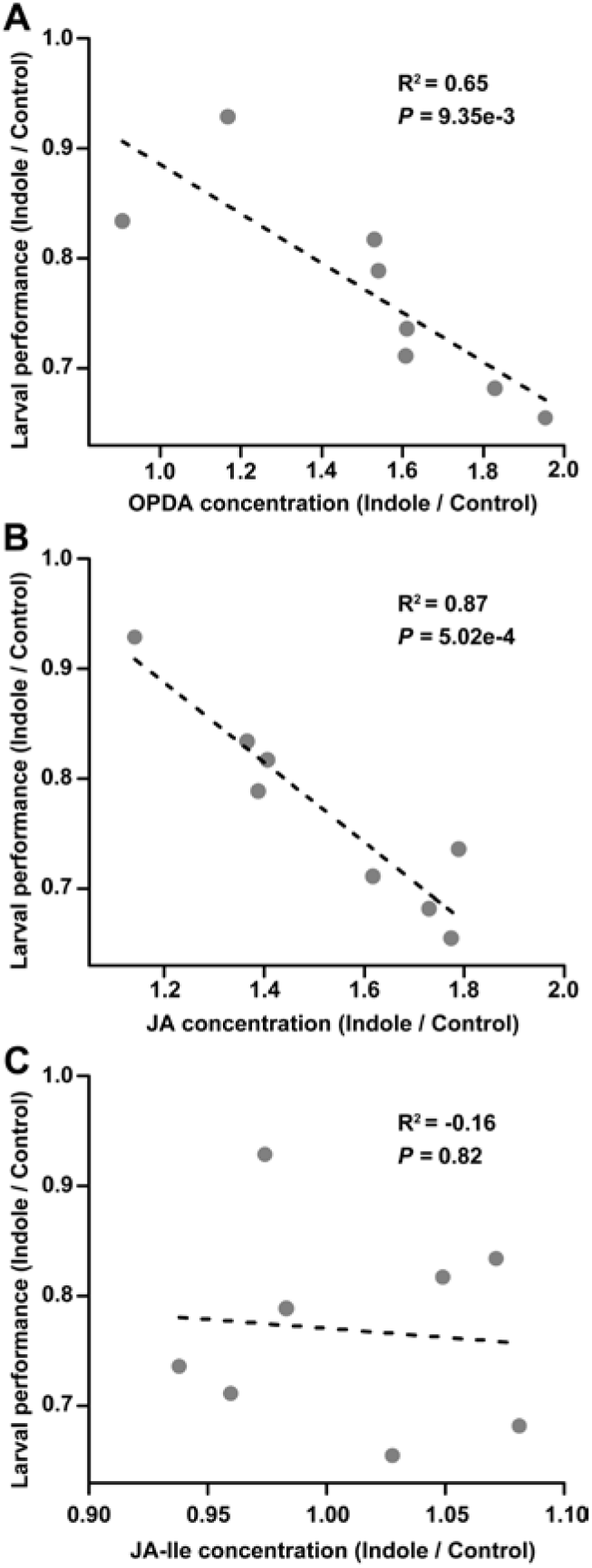
Correlations between indole priming of OPDA, JA and herbivore resistance. (**A – C**) Correlations between the fold changes of herbivore-induced (**A**) 12-oxophytodienoic acid (OPDA), (**B**) jasmonic acid (JA), and (**C**) JA-isoleucine (JA-Ile) concentrations in indole-exposed plants relative to control-exposed plants and fold changes of *S. frugiparda* larval performance on indole-exposed plants relative to control-exposed plants. Circles denote individual genotypes. R^2^ and *P*-values of Pearson Product-Moment correlations are shown.

## Discussion

HIPVs can regulate plant defenses and increase herbivore resistance in many different plant species. However, how volatiles influence early defense signaling, and whether the resulting increase of defense responsiveness increases herbivore resistance, is not well understood. This study contributes to filling these gaps of knowledge by identifying early defense regulators that are involved in volatile defense priming and plant resistance to herbivory.

Indole-exposure at physiological doses resulted in marked changes in the expression of early defense signaling genes. The receptor-like kinase *OsLRR-RLK1* was directly induced, while the MPK *OsMPK3* and the WRKY transcription factor *OsWRKY70* were primed for stronger activation and expression. Experiments with transgenic plants revealed that *OsMPK3* expression is required, and *OsWRKY70* contributes to indole-induced downstream responses. As *OsWRKY70* is regulated by and acts downstream of *OsMPK3* (Li et al., 2015), we infer that indole acts upstream of *OsMPK3*. The fact that the indole-induced priming was not altered in an *OsLRR-RLK1*-silenced line further suggests that the expression of this receptor-like kinase, which can regulate *OsMPK3*, is not directly required for indole-priming. An *OsLRR-RLK1* null mutant would be required to completely rule out the involvement of this gene in indole-dependent downstream responses. In maize, GLV exposure has been shown to directly increase the expression of *MAPK6* and *WRKY12* (Engelberth et al., 2013). In contrast, we find that *OsMPK3* and *OsWRKY70* were not directly induced by indole, and only primed after subsequent herbivore induction. Our recent work in maize confirmed that GLVs directly induce defense genes, while indole primes their expression (Hu et al, under review). Thus, while GLVs and indole both strengthen the jasmonate signaling pathway, their mode of action and integration into early defense signaling is likely different. Priming mechanisms have been elucidated for non-volatile chemical priming agents. In Arabidopsis, β-aminobutyric acid (BABA) acts via the aspartyl-tRNA synthetase IBI1 and induces the expression of a lectin receptor kinase *LecRK-VI.2*, which is in turn required for BABA-induced priming (Singh et al., 2012; Luna et al., 2014). Furthermore, thiadiazole-7-carbothioic acid S-methyl ester (BTH) treatment increases mRNA levels and inactive protein levels of MPK3 and MPK6 which are then activated more strongly upon stress and thereby enhance defense responses (Beckers et al., 2009). Our work shows that naturally occurring volatiles such as indole act by modulating similar components of early defense signaling, but in a different manner. For instance, indole exposure primes MPK activity, but does not directly induce MPK accumulation (Figure 3). It also induces the transcription of a receptor-like kinase, but this does not seem to be required to activate downstream responses. We conclude that indole reprograms early signaling through mechanisms that differ from non-volatile chemical elicitors such as BABA and BTH.

Most HIPVs that enhance defenses have also been shown to prime jasmonate biosynthesis. Indole does the same in maize (Erb et al., 2015) and, as shown here, rice. Our experiments with transgenic plants show that the priming of JA requires *OsMPK3* and is enhanced by *OsWRKY70*, both of which are primed by indole-exposure (Figure 5). We thus infer that JA priming results from the modulation of *OsMPK3*-dependent early defense signaling by volatile indole. As indole-exposure primes JA biosynthesis genes, the capacity of plants to synthesize JA upon herbivore elicitation is likely increased through the higher abundance of rate-limiting enzymes (Haga et al., 2008; Yara et al., 2008; Riemann et al., 2013). *OsAOC*, for instance, which catalyzes allene oxide to OPDA, is encoded by only a single copy gene, and OsAOC-defective rice plants are jasmonate-deficient (Riemann et al., 2013; Lu et al., 2015). Indole exposure also directly induces the accumulation of the JA precursor OPDA. In theory, this bigger pool may increase the formation of JA upon elicitation through the induction of *OsOPR3* following herbivore attack. However, our experiments show that OPDA depletion upon elicitation is not strictly required for JA priming. Thus, there is currently no evidence that direct OPDA induction is causally linked to JA priming in indole-exposed plants.

OsMPK3, OsWRKY70 and JA are part of the same signaling cascade and are positive regulators of rice resistance to chewing herbivores (Zhou et al., 2009; Wang et al., 2013; Li et al., 2015). Indole primes these defense signaling components, and silencing their expression reduces indole-induced resistance against *S. frugiperda*, which illustrates that indole increases plant resistance by enhancing early defense signaling and JA biosynthesis. Thus, apart from repelling and intoxicating certain herbivores (Veyrat et al., 2016), HIPVs suppress herbivore growth and boost plant resistance by enhancing JA-dependent plant defense responses. A recent study documented that pathogen-induced pinenes can trigger systemic acquired resistance (SAR), an effect which was dependent on SA biosynthesis (Riedlmeier et al., 2017). Thus, in analogy to HIPVs, plant volatiles can also trigger resistance against pathogens by enhancing plant defenses through other phytohormonal signaling pathways.

In summary, we propose the following model. Rice leaves that are attacked by herbivores release the volatile indole. Through as yet unknown perception mechanisms, indole primes MPKs in non-attacked tissues. When these tissues come under attack, OsMPK3 is activated more strongly, which boosts downstream responses, including the transcription of *OsWRKY70* and jasmonate biosynthesis genes, which again results in an over accumulation of bioactive oxylipins such as OPDA and JA. Enhanced jasmonate signaling then boosts plant defense responses and thereby reduces herbivore growth and damage. This study provides a mechanistic basis for the regulatory potential and mode of action of HIPVs in plant defense priming.

## Material and Methods

### Plant and insect resources

The rice (*Oryza sativa*) cultivar Xiushui 110 was used in this study. In addition, the transgenic line *ir-lrr1* and its corresponding wild type line Xiushui 110 as well as the transgenic lines ir-mpk3, ir-mpk6, *ir-wrky70, as-aos1* and their corresponding wild type Xiushui 11 were used. These genotypes have been described and characterized previously (Wang et al., 2013; Hu et al., 2015; Li et al., 2015; Hu et al., 2018). Rice seeds were pre-germinated, and then sown in plastic pots (11 cm height, 4 cm diameter) using commercial potting soil (Aussaaterde, Ricoter Erdaufbere-itung AG, Switzerland). Plants were grown in a greenhouse (26°C ± 2°C, 55% relative humidity, 14:10 h light/dark, 50,000 lm m^−2^). Plants were watered three times per week, and used for experiments 30 days after sowing. Fall armyworm (*Spodoptera frugiperda*) larvae were provided by University of Neuchâtel and reared on artificial diet as previously described (Maag et al., 2014). Oral secretions (OS) were collected from third instar *S. frugiperda* larvae which had been feeding on rice leaves for 48 h, and diluted 1:1 with sterilized Milli-Q water before use.

### Quantification of herbivore-induced indole

To determine natural emission rates of indole, we infested rice plants with 3, 5 or 8 third-instar *S. frugiperda* larvae for 12 h, resulting in the consumption of approx. 10%, 30% and 50% of total leaf area. Following infestation, volatiles were collected using a dynamic headspace sampling system and Super-Q traps (n=8). Briefly, the rice plants were enclosed with cooking bags (PET, 35 × 40 cm, max. 200 °C, Migros supermarket, Switzerland). Purified air from a multiple air-delivery system entered the bags via Teflon tubing at a rate of 0.7 L min^−1^ and was pulled out through the Super-Q trap (Volatile Collection Trap LLC., UK) at a rate of 0.3 L min^−1^. Before collection, the Super-Q traps were rinsed with 3 mL of methylene chloride (≥ 99.8 %, GC, Sigma, USA). Volatiles were collected for 8 h. After collection, the traps were extracted with 200 μL of methylene chloride which contains two internal standards (n-octane and nonyl-acetate, each 1 μg in 200 μL methylene chloride). Then, a 1 μL aliquot of each sample was injected into GC/MS (Agilent 7820A GC interfaced with an Agilent 5977E MSD, USA) in pulsed split mode onto an apolar column (HP-5MS, 30 m, 0.25mm ID, 0.25 μm film thickness, Alltech Associates, Inc, USA) for analysis. Helium at constant flow (1 mL min^−1^) was used as carrier gas. After injection, the column temperature was maintained at 40 °C for 1 min, increased to 250 °C at 6 °C min^−1^ followed by a postrun of 3 min at 250 °C. The quadrupole MS was operated in the electron ionization mode at 70 eV, a source temperature of 230 °C, quadrupole temperature of 150 °C, with a continuous scan from *m/z* 50 to 300. The detector signal was processed with HP GC Chemstation software. Absolute emission rates of indole were determined by peak areas, and calculated using a standard curve of synthetic indole (>98%, GC, Sigma, USA).

### Indole exposure

To expose rice to synthetic indole, we covered plants of different genotypes individually with passively ventilated plastic cylinders (40 cm height, 4 cm diameter) made of transparent plastic sheet (Rosco Laboratories Inc., USA). The plants were placed into the greenhouse (26 °C ± 2 °C, 55% relative humidity, 14:10 h light/dark, 50,000 lm m^−2^), and indole or control dispensers were added into the cylinders. After 12 h of exposure, the cylinders were carefully removed and the plants were subjected to OS elicitation (see “Plant elicitation” below). Indole and control dispensers were made as described previously (Erb et al., 2015). Briefly, dispensers consisted of 2 mL amber glass vials (11.6 × 32 mm^−2^; Sigma) containing 20 mg of synthetic indole (>98%, GC, Sigma, USA). The vials were closed with open screw caps that contained a PTFE/rubber septum, which was pierced with a 1 μL micropette (Drummond, Millan SA, Switzerland). The vials were sealed with parafilm and wrapped in aluminum foil for heat-protection and to avoid photodegradation. GC/MS analyses using the approach described above showed that these dispensers release approx. 21 ng h^−1^ volatile indole, which corresponds to amounts emitted by a single rice plant under attack by *S. frugiperda* (Figure 1). Control dispensers consisted of empty glass vials. Dispensers were prepared 24 h before the start of experiments. As we used a passively ventilated cylinder system, indole may accumulate at levels that are higher than expected under natural conditions. To test whether plant defense responses are affected by potential accumulation over time, we exposed rice plants to dispensers for 1 h, 3 h, 6 h and 12 h and measured priming of jasmonic acid (JA) as a downstream defense marker (see sections “plant elicitation” and “phytohormone quantification”). We found that JA priming is independent of the duration of indole exposure (Figure 4). We therefore proceeded in using this system and an exposure time of 12 h for the remaining experiments.

### Plant elicitation

After indole-exposure, cylinders and dispensers were removed. Maize plants were elicited by wounding two leaves over an area (~0.5 cm^−2^) on both sides of the central vein with a razor blade, followed by the application of 10 μL of *S. frugiperda* OS. This treatment results in plant responses similar to real herbivore attack (Erb et al., 2009; Fukumoto et al., 2013; Chuang et al., 2014). Leaves were then harvested at different time intervals, and flash frozen for further analysis.

### Herbivore performance

One starved and pre-weighed second instar larva was individually introduced into cylindrical mesh cages (1 cm height and 5 cm diameter), and clipped on the leaves of rice plants which were pre-exposed to indole or control. The position of the cages was moved every day to provide sufficient food for the larvae. Larval mass was determined 7 days after the start of the experiment. To quantify damage, the remaining leaf pieces were scanned, and the removed leaf area was quantified using Digimizer 4.6.1 (Digimizer) (*n*=15).

### Phytohormone quantification

Rice leaves were harvested at 0, 90 and 360 min after the start of OS elicitation, and ground in liquid nitrogen (*n*>5). The phytohormones OPDA, JA, JA-Ile, SA, and ABA were extracted with ethyl acetate spiked with isotopically labeled standards (1 ng for d_5_-JA, d_6_-ABA, d_6_-SA, and ^13^C_6_ -JA-Ile) and analyzed with UHPLC-MS/MS as described (Glauser et al., 2014).

### Gene expression analysis

Quantitative real time PCR (QRT-PCR) was used to measure the expression levels of different genes. Rice leaves were harvested at 0, 90 and 360 min after the start of OS elicitation, and ground in liquid nitrogen (*n*>4). Total RNA was isolated from rice leaves using the GeneJET Plant RNA Purification Kit (Thermo Scientific, USA). One μg of each total RNA sample was reverse transcribed with SuperScript^®^ II Reverse Transcriptase (Invitrogen, USA) to synthesize cDNA. The QRT-PCR assay was performed on the LightCycler^®^ 96 Instrument (Roche, Switzerland) using the KAPA SYBR FAST qPCR Master Mix (Kapa Biosystems, USA). A linear standard curve was constructed using a serial dilution of cDNA which was pooled from all plants, and generated by plotting the threshold cycle (Ct) against the log10 of the dilution factors. The relative transcript levels of the target genes in samples were determined according to the standard curve. A rice actin gene *OsACTIN* was used as an internal standard to normalize cDNA concentrations. The primers used for QRT-PCR for all tested genes are listed in Supplemental Table 1.

### MPK protein and activation detection

Rice leaves were harvested at 0 and 90 min after the start of OS elicitation, and ground in liquid nitrogen. Total proteins were extracted from pooled leaves of six replicates at each time point using the method described (Wu et al., 2007). Forty μg of total proteins were separated by SDS-PAGE and transferred onto Bio Trace pure nitrocellulose blotting membrane (Bio-Rad, USA). Immunoblotting was performed using the method established previously (Hu et al., 2015). The primary antibody anti-MPK3 (Beijing Protein Innovation, China) or anti-MPK6 (Beijing Protein Innovation, China) was used to detect the total proteins of OsMPK3 or OsMPK6 respectively. The rabbit monoclonal anti-phospho-ERK1/2 (anti-pTEpY) antibody (Cell Signaling Technologies, USA), which is specific for the activated (phosphorylated) form of the p44/42 MPKs (Thr202/Tyr204) (Segui-Simarro et al., 2005; Anderson et al., 2011) was used to detect the active OsMPK3 and OsMPK6. The plant-actin rabbit polyclonal antibody (EarthOx, USA) was used for loading control and detected on a replicate blot. Antigen-antibody complexes were detected with horseradish peroxidase-conjugated anti-rabbit secondary antibody (Thermo Scientific, USA) followed by chemiluminescence detection with Pierce™ ECL Western Blotting Substrate (Thermo Scientific, USA).

### Statistical analyses

Differences in levels of gene expression and phytohormones were analyzed by analysis of variance (ANOVA) followed by pairwise comparisons of Least Squares Means (LSMeans), which were corrected using the False Discovery Rate (FDR) method (Benjamini and Hochberg, 1995). The data normality was verified by inspecting residuals using the “plotresid” function of the R package “RVAideMemoire” (Herve, 2015). The variance homogeneity was tested through Shapiro-Wilk’s tests using the “shapiro.test” function in R. Datasets that did not fit assumptions were log-or asinh-transformed to meet the requirements of normality and equal variance. Differences in larval growth and leaf damage were determined by two sided Student’s t-tests. The relative priming intensity was calculated by the fold changes of larval growth, OPDA or JA levels in the indole-exposed plants relative to control-exposed plants. The differences in fold changes were compared using Student’s t-tests. The correlations (fold changes of OPDA, JA or JA-Ile vs fold changes of larval growth) were tested through Pearson’s product-moment correlation using the “cor. test” function in R (Puth et al., 2014). All the analyses were conducted using R 3.2.2 (R Foundation for Statistical Computing, Vienna, Austria).

### Accession Numbers

Sequence data from this article can be found in the Rice Annotation Project under accession numbers *OsLRR-RLK1* (Os06g47650), *OsHI-RLK2* (Genbank accession number XM_015757324), *OsMPK3* (Os03g17700), *OsMPK6* (Os06g06090), *OsWRKY70* (Os05g39720), *OsWRKY53* (Os05g27730), *OsWRKY45* (Os05g25770), *OsWRKY33* (Os03g33012), *OsWRKY30* (Os08g38990), *OsWRKY24* (Os01g61080), *OsWRKY13* (Os01g54600), *OsHI-LOX* (Os08g39840), *OsAOS1* (Os03g55800), *OsAOC* (Os03g32314), *OsOPR3* (Os08g35740), *OsJAR1* (Os05g50890), and *OsACTIN* (Os03g50885).

## Supplemental Data

**Supplemental Figure 1.**
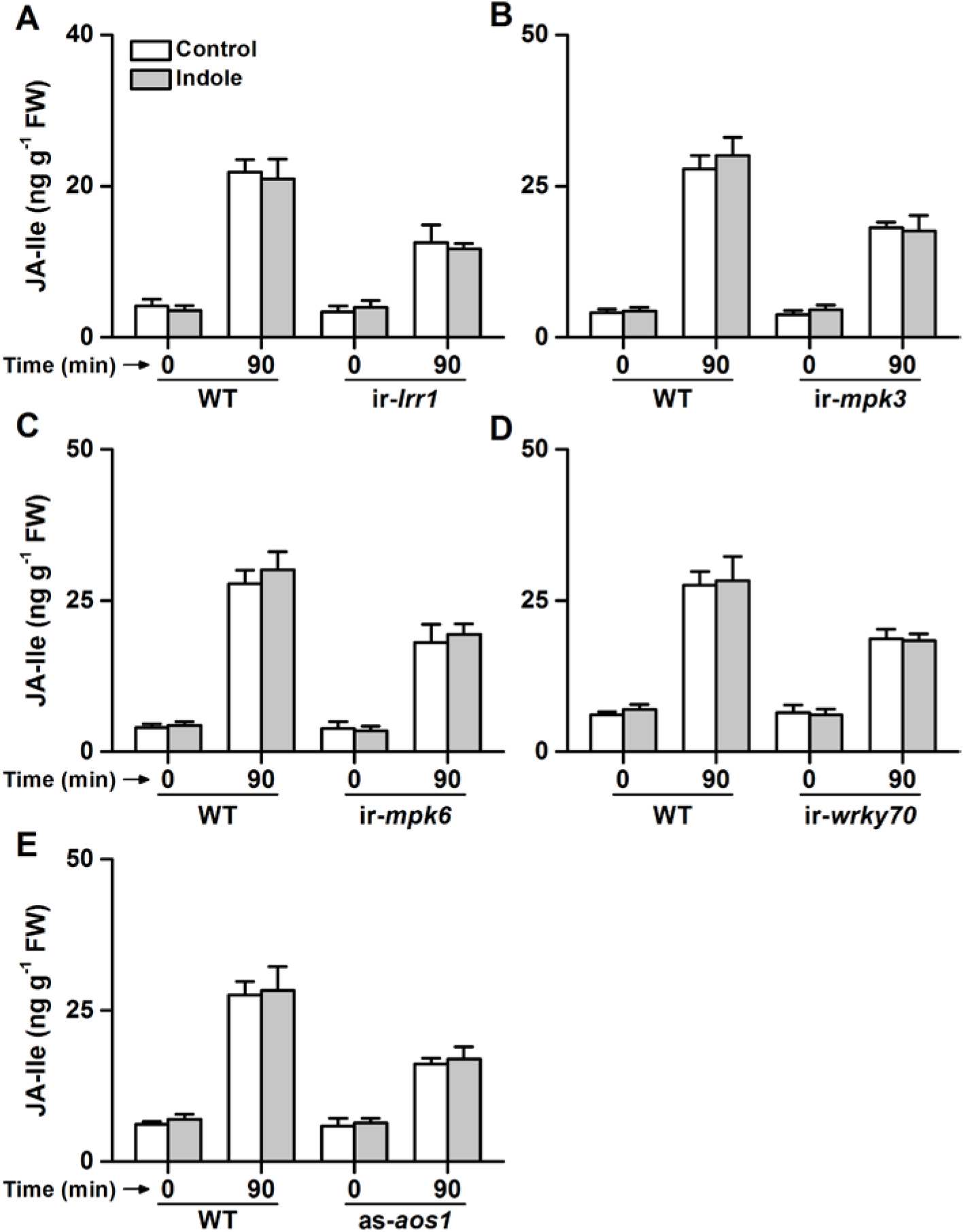
Herbivore-induced jasmonic acid-isoleucine (JA-Ile) levels in MPK, WRKY and JA-impaired plants after indole exposure. Average concentrations of the herbivore-induced JA-Ile in in ir-*lrr1* (A), ir-*mpk3* (B), ir-*mpk6* (C), ir-*wrky70* (D), as-*aos1* (E) line and wild type (WT) plants that were pre-exposed to indole or control (+SE, *n*=6). No significant was found between indole and control treatments at the indicated times. FW, fresh weight.

**Supplemental Table 1.**
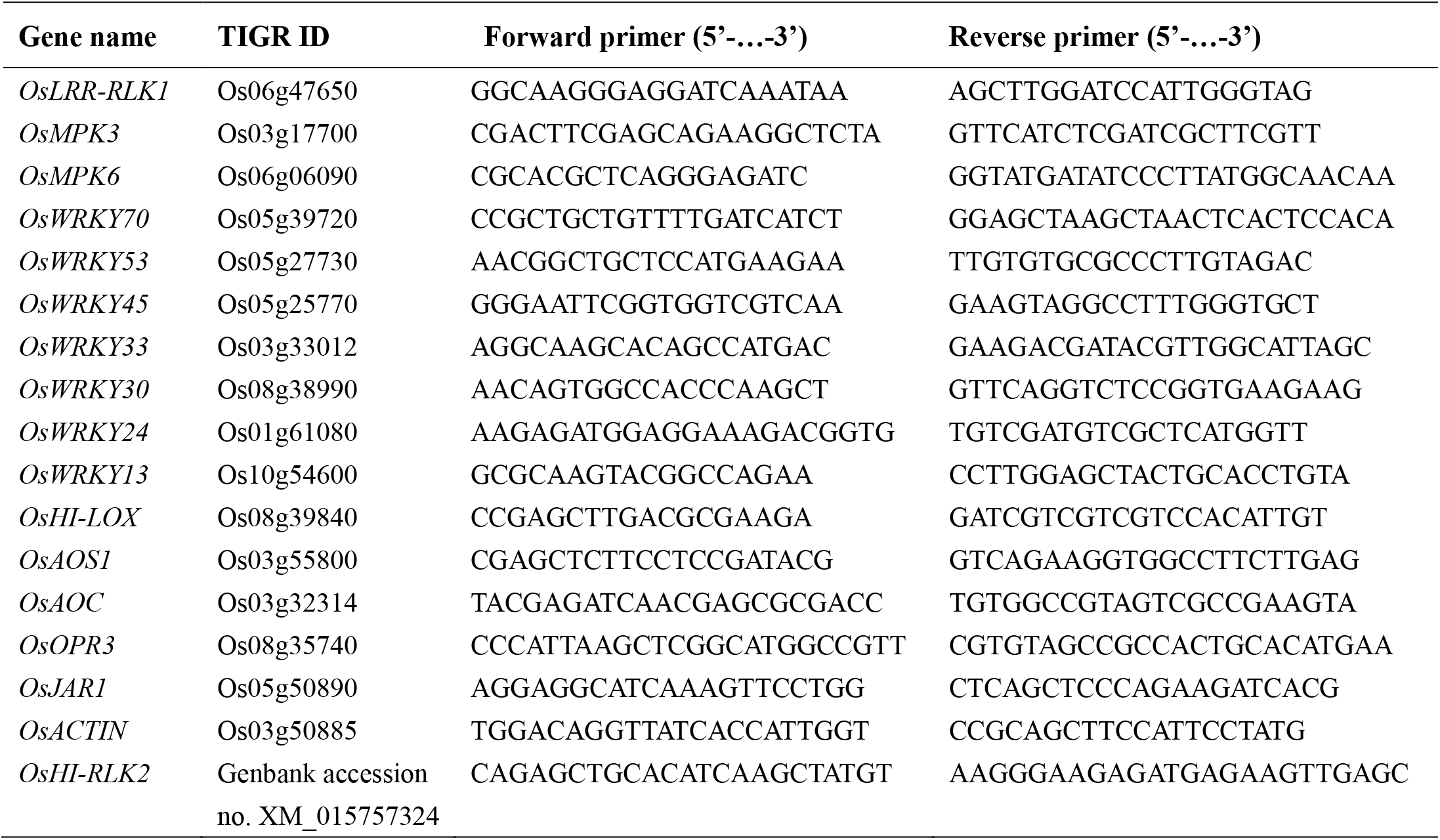
Primers used for QRT-PCR of target genes.

## Acknowledgements

The study was jointly sponsored by the Swiss National Science Foundation (Grant 155781), the Sino-Swiss Science and Technology Cooperation (Exchange Grant Nr. EG 03-032016) and the European Research Council (ERC) under the European Union’s Horizon 2020 research and innovation programme (ERC-2016-STG 714239). The authors declare no conflict of interest.

## Author Contributions

M. E., Y. L. and L. H. conceived the project. M. E and Y. L. acquired project funding. L. H., M. Y., Y. L. and M. E. designed research. L. H., M. Y. and G. G. performed experiments. L. H., M. Y., Y. L. and M. E. analyzed and interpreted data. L. H., M. Y., and M. E. prepared and wrote the first draft. All authors read and approved the manuscript.

